# The interplay between adsorption and aggregation of von Willebrand factor chains in shear flows

**DOI:** 10.1101/2022.12.26.521955

**Authors:** Helman Amaya-Espinosa, Alfredo Alexander-Katz, Camilo Aponte-Santamaría

## Abstract

Von Willebrand factor (VWF) is a giant extracellular glycoprotein that carries out a key adhesive function during primary hemostasis. Upon vascular injury and triggered by the shear of flowing blood, VWF establishes specific interactions with several molecular partners in order to anchor platelets to collagen on the exposed sub-endothelial surface. VWF also interacts with itself to form aggregates that, adsorbed on the surface, provide more anchor sites for the platelets. However, the interplay between elongation and subsequent exposure of cryptic binding sites, self-association, and adsorption on the surface, remained unclear for VWF. In particular, the role of shear flow in these three processes is not well understood. In this study, we address these questions by using Brownian dynamics simulations at a coarse-grained level of resolution. We considered a system consisting of multiple VWF-like self-interacting chains that also interact with a surface under a shear flow. By a systematic analysis, we reveal that chain-chain and chain-surface interactions coexist non-trivially to modulate the spontaneous adsorption of VWF and the posterior immobilization of secondary tethered chains. Accordingly, these interactions tune VWF’s extension and its propensity to form shear-assisted functional adsorbed aggregates. Our data highlights the collective behavior VWF self-interacting chains have when bound to the surface, distinct from that of isolated or flowing chains. Furthermore, we show that the extension and the exposure to solvent have a similar dependence on shear flow, at a VWF-monomer level of resolution. Overall, our results highlight the complex interplay that exists between adsorption, cohesion, and shear forces and its relevance for the adhesive hemostatic function of VWF.

## Introduction

When a vascular injury occurs, the subendothelium of blood vessels gets exposed and in order to stop the bleeding the hemostasis immune response is initiated. During this process, multiple thrombogenic substances interact with each other to create and stabilize a hemostatic plug. Von Willebrand factor (VWF) is among them [1, 2]. VWF anchors platelets to the sub-endothelial surface exposed by the injury[3], triggered by the shear-stress imposed by the flowing blood [4, 5, 6, 7, 8]. Accordingly, malfunction of VWF is related to a broad range of bleeding disorders [9].

VWF is a huge adhesive extracellular-protein biopolymer, with a multimeric structure of variable and exponentially-distributed length [1, 10, 11]. It is composed of multiple covalently-bound monomers, organized in a head-to-head and tail-to-tail sequence [1, 9]. Each mature VWF monomer is made up of 12 protein domains of nanometer size [1]. VWF undergoes reversible conformational changes from a globular to a stretched conformation under a flow [4, 5], thereby exposing binding sites for the interaction with a multitude of partners. Importantly, VWF attaches to the surface by establishing specific interactions with the exposed collagen of the subendothelium, via primarily the VWF A3 domain [12]. Tethered on the surface, VWF immobilizes flowing platelets, via the interaction of its A1 domain with the glycoprotein IB *α* platelet receptor [13, 14]. A reinforcement follows with the binding of the platelet integrin αIIbβ3 to the VWF C4 domain [7, 8].

A key concept is the ability of VWF to interact with itself. Non-covalent self-association of freely-flowing VWF chains on immobilized ones has been previously demonstrated [15, 16, 17]. A consequence of self-association was thus the enhancement of platelet adhesion by providing more binding sites, through both, directly and indirectly, immobilized VWF [18]. VWF has also been observed to bind to platelet-bound VWF, thereby promoting platelet activation [19]. Single-molecule experiments revealed that tension along the tethered chains promoted VWF elongation, which regulates the reversible self-association of VWF [20]. However, there is evidence suggesting that VWF self-associates under static conditions too [21]. Furthermore, multiple protein domains have been found to participate in the self-association of VWF [21], which is also regulated by the A2 domain [22, 23].

Specific domain–domain non-covalent self-interactions are involved in crucial functional aspects of VWF. Examples of this are the force-mediated auto-inhibition of VWF for the binding of platelets mediated by A1–A2 interactions [24, 25, 26, 27], the pH and calcium-dependent stabilization of VWF dimmers imparted by the interaction of the VWF D4 domain [28], and the assembly of VWF tubules for storage, stabilized by D–D and D–A1 interactions [29, 30, 31, 32].

VWF has been the subject of intense studies over the last three decades [1]. In particular, computer simulations have contributed enormously to the functional understanding of this protein. VWF is commonly simulated at a coarse-grained (CG) level of resolution with each CG bead representing a monomer (or a dimer). Such coarsening has enabled simulations to reach relevant spatio-temporal scales and thereby establish the link between the shear-stress imposed by the flowing blood and the elongation propensity of VWF [4]. Simulations also explained the factors governing the formation of reversible polymer-colloid VWF-platelet-like aggregates [6] and the adherence of single-VWF chains to a surface [33, 34, 35, 36], as well as the mechanism underlying enzymatic cleavage [37, 38]. CG simulations also showed that VWF adsorption obeys a cooperative mechanism [39] and added support to the involvement of D4–D4 interactions on the stability of VWF dimers [40].

Despite the wealth of this data, the interplay between three key processes of VWF, namely, elongation and subsequent exposure of cryptic binding sites; chain adsorption, and self-association, in the context of multiple chains subjected to shear flow, remains poorly understood. We addressed this issue by performing Brownian dynamics (BD) simulations of a set of VWF chains under shear flows. We adopted a bead-spring CG representation, in which each chain bead represented a VWF domain. This level of resolution, higher than conventional VWF CG models, allowed us to consider the exact distribution of domains that bind to the surface (namely the VWF A3 domains) and to retrieve more realistic surface area exposures. Our simulations demonstrate a complex interplay in which inter-chain cohesiveness and surface-chain interactions modulate the shear response of VWF chains, tuning their elongation and their propensity to form functional adsorbed aggregates. Our data highlights the different behavior VWF chains have, depending on whether they are bound to the surface or flowing. These findings also show the collective behavior VWF chains exhibit which leads to distinct properties compared to VWF chains in isolation. Finally, we studied the effect the level of resolution has on the chain exposure to the solvent and compare that with the extension, a very common descriptor of the biological function of VWF.

## Methods

### Coarse-grained model

Biopolymers were modeled as chains of interacting beads (Fig. 1A). Neighbor beads (*i, i* + 1) were bonded by harmonic springs, modeled via a potential energy

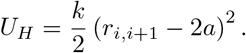

**Figure 1:**
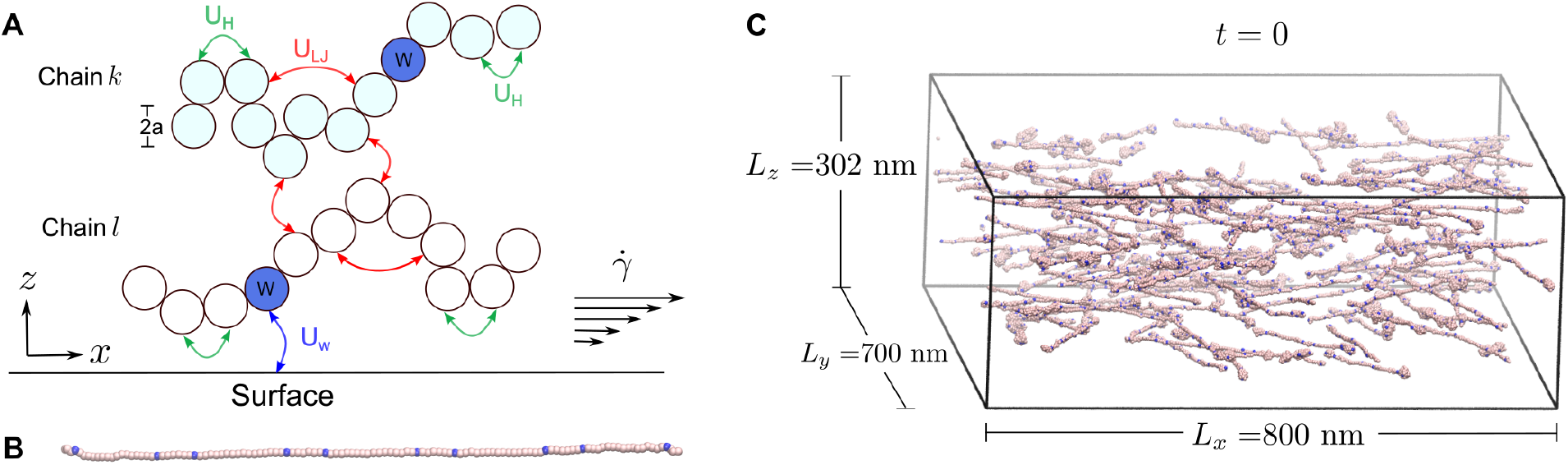
Coarse-grained model to monitor the aggregation and adsorption of VWF-like biopolymers under shear flows. **(A)** The scheme shows two biopolymer chains (here a VWF monomer) composed of several protein domains (circles). The size of one bead resembled the size of the VWF A1 domain (2*a* = 3.2 nm). Neighbor beads interact via bonded harmonic interactions *U_H_*. Non-neighbor bead interactions were modeled through a short-range Lennard-Jones potential *U_LJ_*, both in (here chain *k*) or across chains (*k* and *l*). The potential *U_w_* takes into account the interaction of the beads with the surface. A particularly strong interaction for the bead corresponding to the VWF A3 domain (blue W-labeled bead) was considered. A shear flow with a shear rate 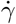 was imposed on the chains as indicated by the arrows. **(B)** Biopolymer chains of *n*=120 domains, corresponding to a VWF decamer, were considered. Here a fully extended conformation of one of these chains is shown. Domains specifically interacting with the surface were distributed along the chain based on the positions of the VWF A3 domains [1] (labeled here as W and depicted as blue spheres). **(C)** Initial configuration (*t* = 0) of the simulated system consisting of 200 biopolymer chains, with different extensions and randomly-distributed positions and orientations. Simulation box dimensions are indicated.

Here, *r*_*i,i*+1_ is the distance between consecutive beads, *a* the bead radius, and *k* the spring constant (a soft spring *k* = 100 kJ/mol·nm^2^ was used). Volume exclusion between pairs of beads (*i, j*) was considered through a shortrange Lennard-Jones potential

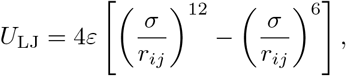

with *r_i,j_* the distance between beads *i* and *j. σ* was set to 2a, i.e. related to the minimum equilibrium separation between beads. The strength of the non-bonded interaction e was normalized by the thermal energy, *k_B_T*, with *k_B_* the Boltzmann constant and *T* the temperature (*k_B_T* ~2.5 kJ/mol at 300 K). Accordingly, normalized values 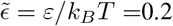, 0.4, 0.6, 0.8, 1.0 were considered. This range allowed the chains to be extended by shear flows [4].

Interactions with the bottom and top surfaces of the simulation box were modeled by a 10-4 potential [41]:

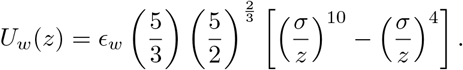

Here, *z* is the distance from the bead to either surface. The strength of the interaction was controlled with the parameter *ϵ_w_*. This parameter was fixed at a weak value at the top surface (*ε_w_* = 10^−5^kJ/mol, i.e. 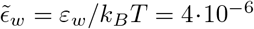) to prevent accumulation of chains on this surface. The adsorption of chains at the bottom surface was of interest and therefore the interaction energy with that surface was increased 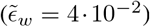. Note that this value was varied for a specific set of beads resembling the VWF A3 domains (see below). The same *σ* value was used here as with the Lennard-Jones potential used between bead pairs.

To mimic VWF biopolymers, each bead was assumed to represent one VWF protein domain (i.e. *a* = 1.6 nm according to the radius of gyration obtained for the VWF A1 domain by molecular dynamics simulations [26]). In addition, beads corresponding to the VWF A3 domain (labeled here as W in Fig.s 1A, B) were assumed to have an increased and varying interaction energy with the surface 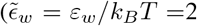, 6, and 10), according to previously-chosen ranges [42]. We considered chains of *n*=120 beads, corresponding to 10 mature VWF monomers, each one of them composed of 12 protein domains [1] (Fig. 1B).

The receptor density on the surface, *φ*, was set *φ* = 10^−2^ nm^−2^, which corresponds roughly to 1 receptor in a squared area covered by one globular VWF monomer. See the results below for a justification of this density choice.

### Brownian dynamics

Biopolymer dynamics was simulated with conventional Brownian Dynamics (BD) simulations. The velocity *v_i_* of the i-th bead was described by the Langevin equation [43, 4]:

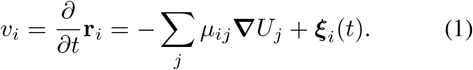

The first term on the right-hand side represents the velocity component associated with diffusion, which depends on the total force acting on the bead (***F**_j_* = -**▽***U_j_*) and the mobility tensor *μ_ij_*. The term *ξ_i_* is related to the random forces exerted by the solvent molecules over each bead. *ξ_i_*(*t*) satisfies the correlation function 〈*ξ_i_*(*t*)*ξ_j_*(*t’*)〉 = 2*k_B_Tμ*_0_*δ_ij_δ*(*t – t’*), with *μ*_0_ being the mobility of each bead. In the free draining regime (no hydrodynamic interactions between beads, i.e. *μ_ij_* = *μ*_0_*δ_ij_*), the equation (1) can be solved numerically at discrete time steps Δ*t* to obtain a recursive formula for the bead positions ***r**_i_* in the presence of a shear flow [43]:

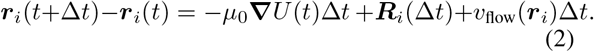

The mobility was obtained from the diffusion coefficient *μ*_0_ = *D*_0_/*k_B_T. D*_0_ was assumed to be 1.32 · 10^−4^ nm^2^/ps, an estimate for the VWF A1 domain translational diffusion coefficient, obtained from molecular dynamics simulations [26]. ***R**_i_*(Δ*t*) is a random number drawn from a Gaussian distribution with zero mean and width 2Δ*tD*_0_. *v*_flow_ is an additional velocity term to take into account the shear flow: 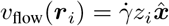, with *z_i_* the position of each bead along the *z* coordinate, and 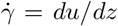 the shear rate, i.e. the gradient of the flow velocity (*u*) along the *z* axis. Note that this third term only affects the *x* component of the bead velocity (Fig. 1*A*). We expressed the time-scales in terms of the characteristic diffusion time of a single bead *τ* = *a*^2^/*D*_0_ = 1.939 · 10^4^ ps. Accordingly, five dimensionless shear rates were considered: 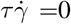, 0.1, 0.25, 0.67, and 1.55.

### Simulation details

Brownian dynamics simulations were carried out with the particle-dynamics simulation toolkit GROMACS [44] (2020.1 version). The velocity term corresponding to the shear flow described in equation (2) was included by modifying the Brownian dynamics position iteration formula in the GROMACS source code. 200 chains were considered. Initial conformations for the chains with different extensions were considered. The 200 chains were sequentially inserted at random positions within a simulation box of dimensions *L_x_* = 800 nm, *L_y_* = 700 nm, and *L_z_* = 302 nm (Fig. 1C). The box boundary conditions were defined as periodic at the *x* and *y* axes, and reflective at the *z* axis. The box dimensions were big enough to accommodate each biopolymer chain in a fully stretched conformation. To remove steric clashes, a steepest descent energy-minimization was carried out before the Brownian dynamics simulations. Trajectories were obtained by numerically integrating equation (2) at discrete time steps Δ*t* = 1 ps (~ 5.16 · 10^−5^*τ*) over a total simulation time of of 500 *μ*s (~ 2.58 · 10^4^*τ*). Unless stated otherwise, the first 0.1 *μ*s (~ 5.16 *τ*) were accounted as equilibration time and thus discarded from the analysis. In total, 75 trajectories were generated by systematically varying 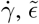 and 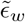.

For the purpose of comparing the dynamics, a set of simulations considering only a single chain, either freely flowing or interacting with the surface were carried out, using identical simulation parameters.

### Simulation analysis

The following observables of the chains were extracted from the simulations, as a function of the dimensionless shear rate 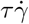, the inter-chain cohesion 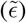, and the chain adsorption 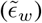 energies.

### Spatial distribution

To identify bulk and adsorption regions, the distribution of the chains along the normalized coordinate *Z* = *z/L_z_* was evaluated (Fig. S1). Chain center of mass *Z* positions (*Z*_COM_) were used for this purpose.

### Extension

The radius of gyration (*R_g_*) was considered as a descriptor of the extension of the chains. The time-averaged square radius of gyration 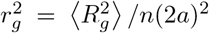, normalized by the mean square end-to-end extension of an ideal chain *n*(2*a*)^2^, was computed for each chain (〈〉 denotes average). Subsequently, the 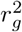 values of all chains were averaged in order to get a global measure of the extension, i.e. 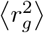. In addition, the ratio 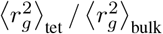 was computed to contrast the extension of the chains that are adhered to the surface (numerator) with respect to the freely flowing bulk ones (denominator).

### Adsorption and tethering

A criterion based on the chain center of mass position is not sufficient to determine if a chain is attached to the surface or not. A more strict attaching criterion was based on the position of each bead. If a bead was at a height *z* ≤ 2*σ*, then the chain this bead belonged to was assumed to be attached to the surface (from here on called an “adsorbed” chain). In addition adjacent chains that were in contact with the absorbed chains (distance ≤ 2*σ*), but not in direct contact with the surface were also monitored (denoted as “adjacent” chains). We then considered the total number of chains tethered to the surface as the sum *N*_tet_ = *N*_ads_ + *N*_adj_, including both the number of adsorbed (*N*_ads_) and adjacent (*N*_adj_) chains. The mean 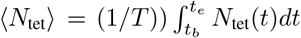 was used as descriptor of level of adsorption, with *T* = *t_e_* – *t_b_* the total duration of the simulation (*t_e_* ~ 2.58 · 10^4^*τ*) minus the initial equilibration time (*t_b_* ~ 5.16 *τ*).

### Aggregation

We used the solvent-accessible surface area [45] to quantify chain aggregation. The value *S* = *S_T_*/Σ*_k_S_k_* is the ratio of the total exposed surface area of the chain aggregate *S_T_* to the sum of the areas *S*_k_ of each chain. Accordingly, a value *S* of 1 indicates that chains were fully dissociated, while a *S* value near to 0 implies they are associated to form a condensate. Similar to the other observables, *S* was evaluated separately for adsorbed (*z* ≤ 2*σ*) and adjacent chains, but also for bulk chains (2*σ* < *z* ≤ *L_z_*).

### Surface area exposure

The total exposed area *S*_T_ of the tethered chains was also normalized by the number of tethered chains *N*_tet_, *S*_c_ = *S_T_/N_tet_*, to obtain the amount of exposed area of each individual chain. Time averages 〈*S*_c_〉_tet_ were computed. 〈*S*_c_〉_tet_ was computed for two different CG resolutions: one bead representing either one VWF protein domain or a VWF monomer. For the latter, the BD trajectories (at the protein-domain level of resolution) were mapped to a monomer-level of resolution by computing the trajectory of the center of masses of the monomers (10 monomers for each chain). The monomer bead radius was estimated to be 10 nm, according to the radial distribution function of the center of masses of the monomers (see results below). Either the protein domain size (a) or the VWF monomer dimension (10 nm) were used as a probe radius for the calculation of *S_T_* and *S*_c_. To compare the exposures to the chain extension, both exposure and extension were normalized by the maxima 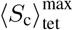 and 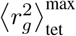, respectively.

### Error estimation

Standard errors were obtained as 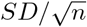, with *SD* the standard deviation of the computed quantities and *n* the number of data samples. *n* was either the number *n_t_* of uncorrelated samples from the time series or the number *n_s_* of chains for the calculation of the chain extension. In the case of the number of adsorbed chains, bootstrapping was used to estimate the error. Accordingly, the time *T* was divided into 5000 segments. The segments were combined randomly allowing replacement, to obtain 〈*N*_ret_〉 in ten runs. The error was estimated as the standard deviation of these ten values.

## Results

We used Brownian dynamics simulations to monitor the dynamics of linear biopolymers in the presence of an external shear flow. The biopolymers were modeled as linear bead-spring chains and mimicked the behavior of the multidomain protein von Willebrand factor.

In our analysis, we distinguished chains that were freely flowing from those that were adsorbed and tethered to the surface. We quantified the distribution of the chain center of mass positions along the *Z* coordinate (perpendicular to the direction of flow) (Fig. S1). Based on this histogram we defined the height *Z*_cutoff_ ~ 7.12 × 10^−2^, normalized by the box size (which is equivalent to 6.72 *σ*) as the cutoff to define tethered (*Z* ≤ *Z*_cutoff_) and bulk (*Z* > *Z*_cutoff_) regions. Accordingly, at that height, the interaction of chains with the surface is negligible, *U_w_* (6.72*σ*) = −1.5 × 10^−3^*ϵ_w_*.

Figure 2 shows examples of final conformations obtained by Brownian dynamics simulations in two typical situations, e.g. in the absence of (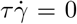, Fig. 2A) or in the presence of a moderate flow (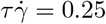, Fig. 2B), respectively. In the absence of flow, the chains adopted a compact configuration. On the contrary, as a consequence of the flow, the shear stress overcame the cohesion forces and the chains stretched along the shear direction rather independently of their initial degree of extension. In addition, chains were adsorbed onto the surface. Furthermore, they formed flowing or tethered conglomerates. In the following sections, we systematically analyze these three different processes, i.e. extension, adsorption, and aggregation, their dependence on the shear flow, and their relevance for the function of the Von Willebrand factor.

**Figure 2:**
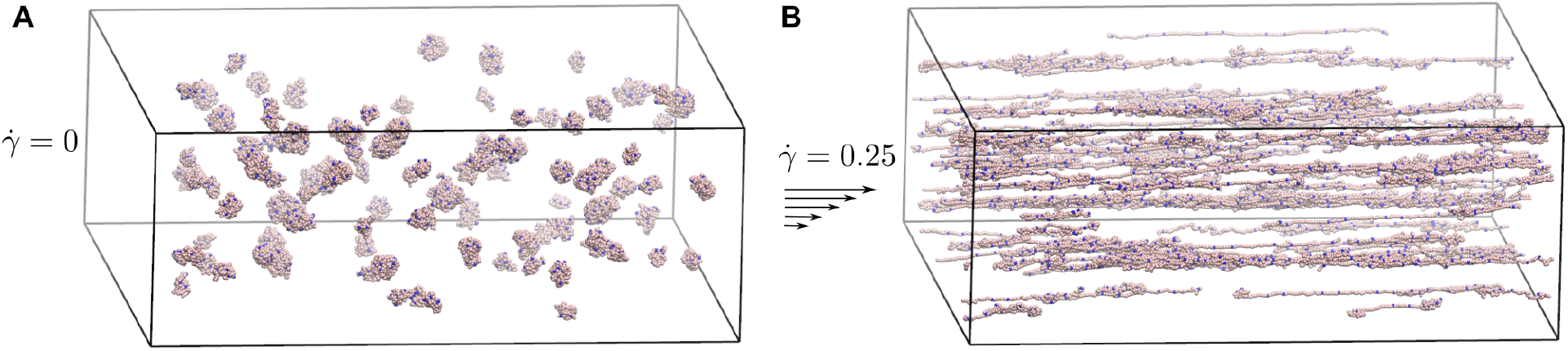
Brownian dynamics simulations of multi-chain VWF-like biopolymers. Examples of the final conformations at two typical conditions, at equilibrium conditions in the absence of flow, 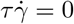 (A: compact conformations) or in the presence of a moderate shear rate 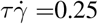 (B: stretched conformations).

### Extension

First, we validated our simulation protocol by comparing the dynamics of single chains with previous theoretical and simulation estimates. In the absence of flow, chains in groups displayed an extension 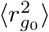 in the range from 0.152–0.182 (Fig. S2A), while the extension of a single isolated chain, 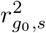, varied from 0.150 to 0.264 (Fig. S3). These values are similar to the extension of an ideal chain [46], i.e. 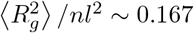. The flow dramatically increased the extension, both of a single chain or a concentrate of them (Fig. 3A–B). In particular, for a single bulk chain with a cohesion strength of 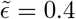 and in the presence of a flow of 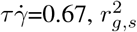 was found to be ~ 6 (Fig. 3A). This means 〈*R*〉 /(2*na*) ~ 0.533, a normalized extension that compares well with previous estimates for similar shear and cohesion values (~ 0.54–0.56) [4].

**Figure 3:**
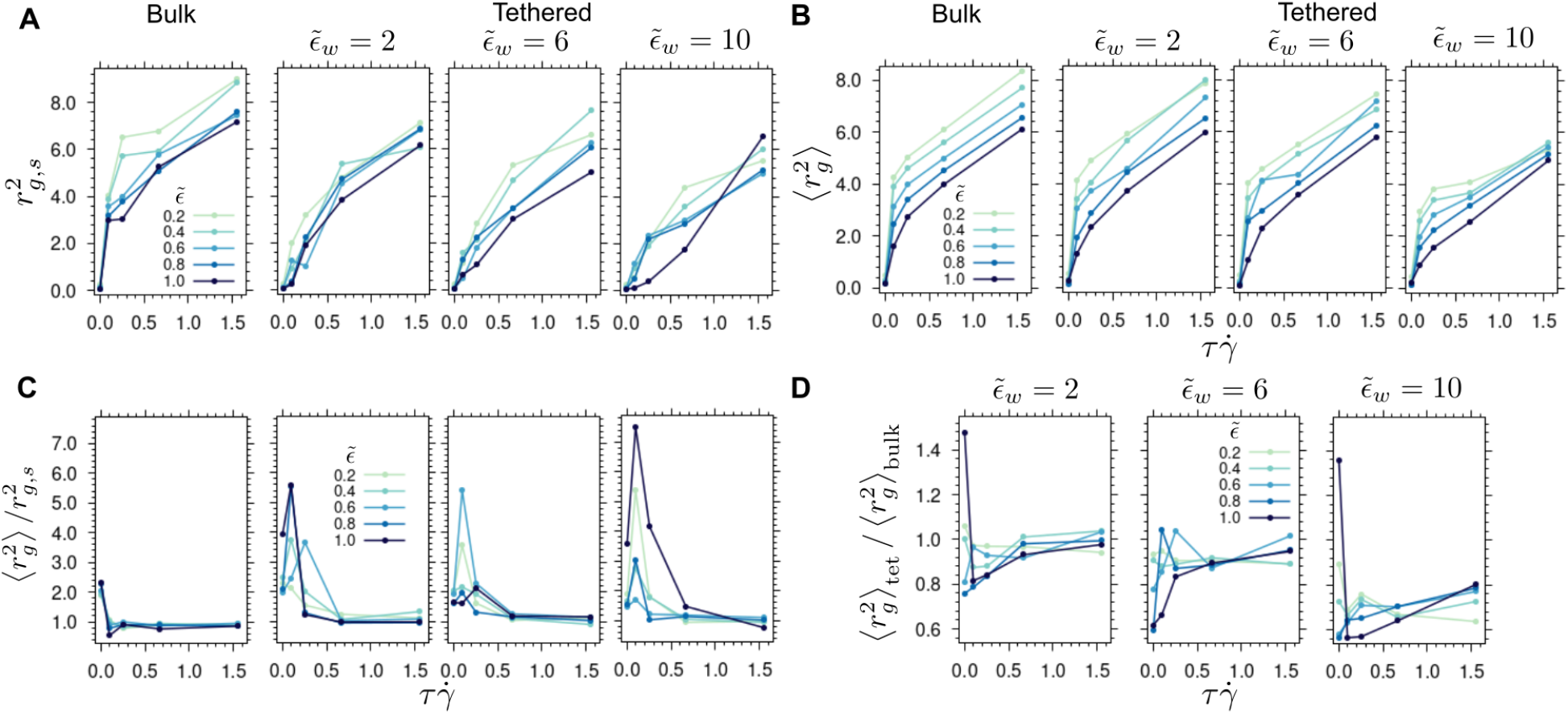
Extension of VWF-like biopolymers under shear flow. (A–B) Normalized mean square radius of gyration 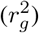 for a single chain in isolation (A) or in a multi-chain system (B) as a function of the shear rate 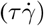, for different polymer–polymer cohesion 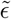 (color) and polymer–surface 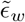 (panels) energies. The calculation was carried out for both bulk (left panel) and tethered (two middle and right panels) regions. 〈〉 denotes the average over chains. Note that in the bulk case, simulations with different 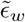 yielded almost identical results (Fig. S4) and thus were averaged into a single curve. The maximum standard error in A was ~ 2.64 and in B it was ~ 0.554 (see errors in detail in Fig. S5). (C) Ratio between the average extension of a chain in the system with multiple chains and that of one chain in isolation, 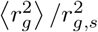 (same format as in A and B). (D) Ratio between the extension of tethered chains and that of bulk chains in the multi-chain system: 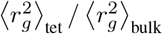 (same format as in A and B).

Next, we compared the dynamics of a single chain (Fig. 3A) with that of a group of chains (Fig. 3B). As expected for the explored ranges of rates (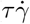 from 0.1–1.5) and chain cohesion strengths (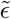 from 0.2–1.0), a monotonic increment in the extension was observed when augmenting the shear flow [4]. The cohesion energy shifted down this increase, although less notoriously for single chains than for interacting ones (compare Fig.s 3A–B). To better quantify the collective behavior of the chains, we monitored the ratio between the chain extension for the multi-chain and the single-chain systems, 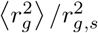 in Fig. 3C. In the presence of flow, bulk chains were found to elongate almost equally, regardless of whether they were in isolation or in groups (values of around 1 for 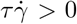 in Fig. 3C, left panel). On the contrary, at shear rate zero, inter-chain interactions promoted a more extended conformation of the bulk interacting chains (see ratio larger than 1 for 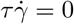 in Fig. 3C, left panel). Interestingly, for groups of chains tethered to the surface, the extension was more enhanced compared to the extension of an isolated chain, and this time not only without the shear but also for other small shear-rate values (Fig. 3C, two middle and right panels). As a consequence, in groups, VWF-like biopolymer chains display a different flow-induced elongation propensity, compared to when they are in isolation.

We also checked the impact tethering has on the extension of the interacting chains. For this, we assessed the extension of the tethered chains with respect to the freely bulk flowing ones (Fig. 3D). For low and medium surface-chain interaction strengths, tethered chains stretched to the same extent as the bulk flowing ones (ratios around 1 for 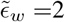 and 6 in Fig. 3D), although with more variability, due to the chain cohesiveness, for zero shear or low-shear values (compare variations in the ratio for low, 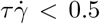, and large fluxes, 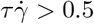 in Fig. 3D). On the other hand, for a high chain-surface interaction 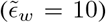, the tethered chains were observed to stretch significantly less than their bulk counterparts, pretty much independently of the internal chain cohesion strength (see the right panel in Fig. 3D and Fig. S2B). Thus, chain cohesiveness together with surface adhesiveness non-trivially modulate the extension of groups of chains.

### Adsorption and Tethering

We next investigated the spontaneous adsorption and posterior tethering of the chains. Fig. 4A shows the number of tethered chains as a function of time for all 75 simulations we carried out, which drastically varied depending on the applied flow, and the chain–chain cohesion and chain–surface interaction energies. We considered the mean 〈*N*_tet_〉 to quantify the overall adhesion to the surface. As expected, the number of tethered chains increased as the chain–surface interaction energy augmented (Fig. 4B). For each of such energies, the shear flow caused a reduction in 〈*N*_tet_〉 (Fig. 4B). No clear trend with the cohesion energy 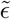 was observed, although it is interesting to note that a large variability between the curves for the different cohesion values (variations in 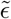) was observed as the surface adhesion energy 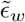 increased (Fig. 4B).

**Figure 4:**
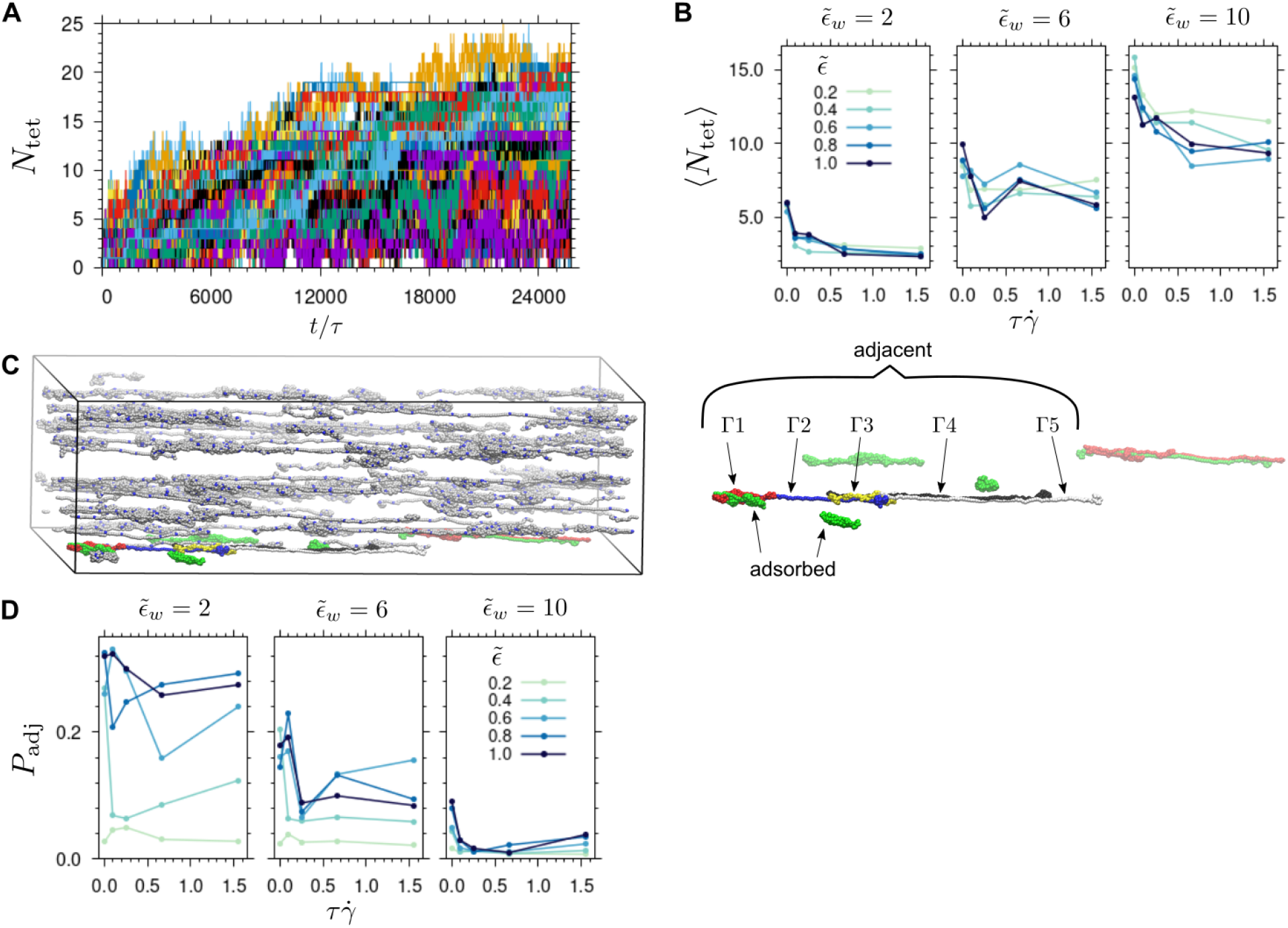
Adsorption and tethering of VWF-like biopolymers under shear flow recovered from BD simulations. (A) The number *N*_tet_ of chains tethered to the surface is displayed as a function of time (in *τ* units) for all 75 carried-out simulations. *N*_tet_ = *N*_ads_ + *N*_adj_, including the number of adsorbed chains *N*_ads_ and of adjacently immobilized chains *N*_adj_, i.e. the chains that were in contact with the adsorbed chains but not in direct contact with the surface. (B) Mean of *N*_tet_ as function of the shear flow 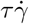, and the polymer–polymer cohesion 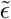 (color) and the polymer-surface 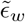 interaction (panels). The maximum value of the error in 〈*N*_tet_〉 was estimated by bootstrapping to be 0.1 (Fig. S6A). (C) The snapshot highlights the tethered chains (color) and the bulk ones (gray), and indicates the different types (Γ from 1-5) of adjacent chains, sequentially tethered to one adsorbed chain. (D) Probability *P*_adj_ of observing at least one adjacent chain attached to an adsorbed chain as a function of the shear flow 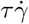, the chain-chain cohesion 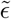 (color) and the chain-surface interaction 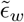 (panels) (same format as in B).

Interestingly, adsorbed chains also formed aggregates with other adjacent chains which were not in direct contact with the surface. Accordingly, sequences of secondary chains tethered to the surface were established (Fig. 4C). States with multiple adjacent chains were rather short-lived and they mainly consisted of the second layer of neighbors(Fig. S7). The probability *P*_adj_ of observing at least one adjacent chain was especially high, and *P*_adj_ increased as cohesion augmented, for moderate chain–surface interaction energies and flows (Fig. 4D).

Thus, the chain–surface energy and the strength of the shear flow largely dominate the amount of tethered chains. In turn, cohesion between chains plays a minor but still noticeable distorting role, contributing to the cooperative tethering of secondary chains.

### Aggregation

We continued our study by analyzing the way chains aggregated with each other. As a measure of aggregation, we used the quantity *S* which is defined as the solvent-accessible surface area of all polymer chains together, divided by the sum of the areas of the individual chains (see methods). The highest possible aggregation was obtained in the absence of flow (see lowest values of *S* below one in figure 5). Augmenting the flow, mostly diminished the level aggregation (increase 〈*S*〉 to values near one) in a hyperbolic fashion. This trend was tuned by the cohesion energy 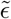, rather independently of whether the chains were bulk or tethered ones (Fig. 5).

**Figure 5:**
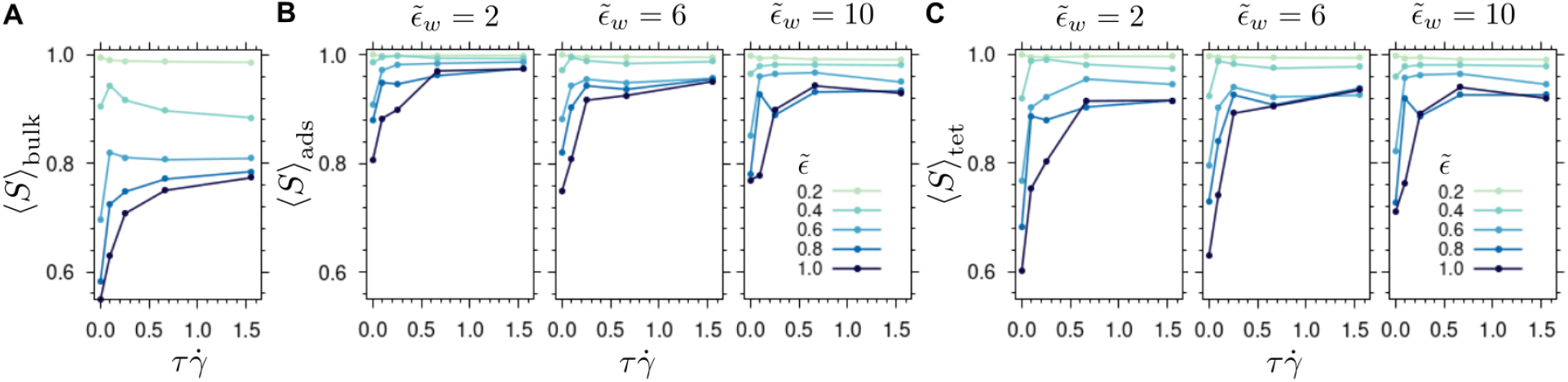
Aggregation of biopolymer VWF-like chains recovered from the simulations. Time-averaged weighted surface exposure ratio, 〈*S*〉, for bulk (A), adsorbed (B), and tethered chains (C) is presented as a function of the shear rate 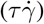, the polymer cohesion 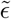 (color) and polymer–surface interaction strength 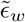 (panels). Note that 〈*S*〉_bulk_ was found to be practically invariant to changes in 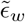 (Fig. S8). Accordingly, here, the average between the values obtained for three different energies 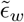 is shown. The maximum standard error of *S* was 0.039 (A), 0.146 (B), and 0.127 (C)(see errors in detail in Fig. S9).

Together, cohesion and adsorption strengths induced intriguing behaviors described as follows. Bulk chains were largely influenced by the cohesion energy 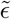 (Fig. 5A), displaying a non-monotonic reducing aggregation dependence on the flow for intermediate cohesion values (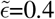 and 0.6), while increasing monotonically for high values (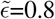 and 1.0). Adsorbed chains were less susceptible to changes in the cohesion energy and showed less aggregation than the bulk ones (compare Fig.s 5A and 5B). On the other hand, when adjacent chains were also considered, a stronger aggregation response was observed at low flows (compare Fig.s 5B and 5C). This constitutes another indication of the previously-shown, interaction of adjacent chains with the adsorbed ones (see Fig.s 4C–D). Nonetheless, both adsorbed and tethered chains reached a similar aggregation degree for the high considered shear rates (see 〈*S*〉 for 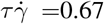 and 1.55 in Fig.s 5B–C), indicating that the cooperative coexistence of tethered aggregates occurred principally at low values of flow. Interestingly, the interaction energy with the surface had a rather moderate influence on the adsorbed chains (i.e. in direct contact with the surface) but a notable diminishing effect in the aggregation of the tethered ones (i.e. including the adjacent chains too) (compare distinct 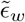 in Fig.s 5B–C).

In summary, aggregation is largely modulated by the cohesion between chains and the shear flow. The interaction of the chains with the surface also influences this process, by altering the aggregation of chains that are not in direct contact with the surface.

### Receptor density

The receptor density on the surface, *φ*, is unknown. The collagen I surface density of 7.5· 10^−4^ molecules/nm^2^ [47] may be considered as a lower boundary because each collagen I molecule could provide multiple receptor sites. In our simulations, we set *φ* = 10^−2^ nm^−2^, which corresponds roughly to one receptor in a squared area covered by one globular VWF monomer (given that the radius of a VWF monomer is ~10 nm, see below). We considered this density to sample a sufficient number of adsorption events while avoiding the formation of an exclusion region. (Fig S10). Note that the extension and aggregation of the bound chains did not drastically change when decreasing the density by a factor of ten, closer to the density used in a previous study of about 1/8 receptors per monomer area [39] (Fig. S11).

### Chain surface exposure

The extension of the chains has been typically considered as the main descriptor of VWF shear-response. However, the protein surface area exposed to the solvent is a quantity that is closer to the main physiological action of the flow on VWF, namely, exposing cryptic binding sites to enable the interaction of VWF with its partners, such as collagen or platelets. We thus took advantage of the surface area calculations to investigate the actual level of exposure of the VWF chains.

At a level of resolution of one bead representing a protein domain, the recovered surface exposure area of all tethered chains together 〈*S*_c_〉_tet_ varied from 2×10^3^ nm^2^ (in the absence of flow) to areas of roughly 7×10^3^ nm^2^ (Fig. 6A). By mapping the conformations into a lower resolution model, in which one bead represents a monomer (i.e. 12 protein-domain beads), exposure values were up to five-fold higher (Fig. 6B). At both resolutions, the surface area increased with the flow and decreased with the cohesion energy in a monotonic fashion (Fig. 6A–B). However, at the monomer level of resolution, a stronger dependence on 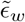 was obtained (Fig. 6B). We compared the exposed surface area with the extension of Fig. 3B (two middle and right panels) in figures 6C–D. Exposed surface areas grew suddenly as the flow increased, i.e. high degrees of exposure were already obtained at small flow values, while the extension augmented more gradually with the flow (Fig.s. 6C-D). Accordingly, the correlation between the exposed area and the extension is not trivial. Remarkably, the exposure at the domain level is almost non-sensitive to the chain-surface interaction energy (Fig. 6C). In contrast, at the monomer level, it exhibits a similar attenuation with this energy as the extension (Fig. 6D). Thus the adopted level of resolution impacts the surface area chains expose in dependency on the surface adhesion energy.

**Figure 6:**
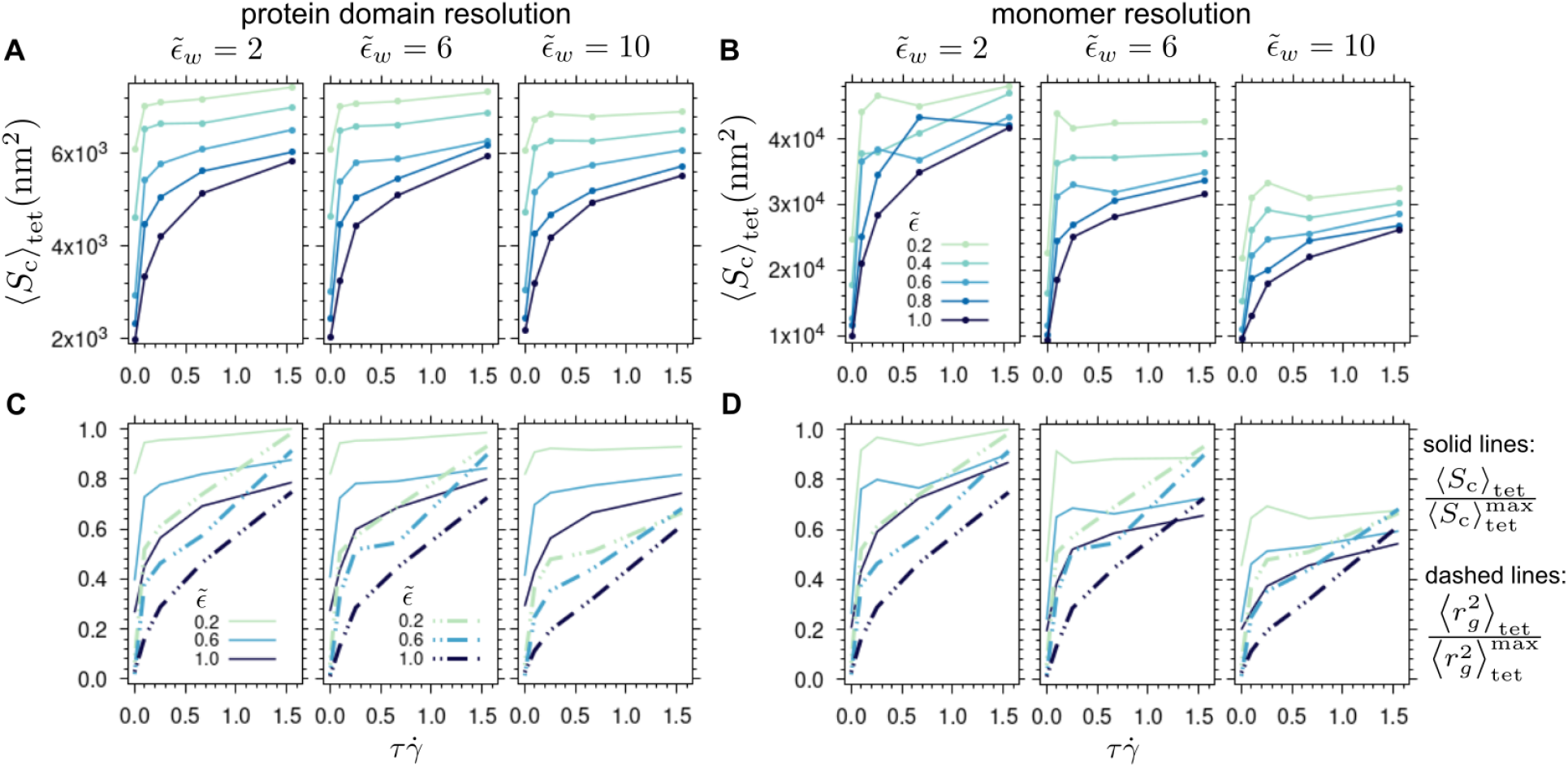
Surface exposure of biopolymer VWF-like chains. (A–B) Time-averaged surface exposure of all tethered chains together (〈*S*_s_〉_tet_) was computed at a level of resolution of one bead representing a protein domain (A) and a bead corresponding to a monomer, i.e. 12 beads (B). Exposure was computed as function of the shear rate 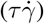, the chain-chain cohesion 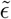 (color), and chain-surface interaction 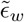 (panels) strengths. (C–D) Normalized exposures (solid lines from A and B) are compared to normalized chain extensions (dashed lines, extracted from Fig. 3B, two middle and right panels) for both levels of resolution. The highest observed value from all the simulations was considered as the normalization factor. The same format is used as in A and B. Plots for 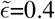 and 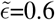 were omitted for clarity. The maximum value of the standard error in *S*_c_ was 903 nm^2^ (see errors in detail in Fig. S6B).

## Discussion

Here, we used Brownian dynamics simulations to systematically examine the elongation, adsorption, aggregation, and surface exposure of biopolymers that resembled the extracellular blood protein VWF under shear flows.

By means of a parameter exploration, we retrieved different regimes in which intra- or inter-chain cohesion, together with the adhesiveness to the surface, modulated the response to shear flow. Overall, there is no dominant factor, but it is the coexistence between them that dictates the chain behavior. For instance, the extension was not only influenced by the cohesion between chains but also by the interaction of them with the surface, i.e. the stronger the latter the more compact chains were (Fig. 3B). Reciprocally, the adsorption of chains was mainly modulated by their interaction with the surface but the cohesive interactions between beads distorted this trend (Fig. 4B), promoting secondary tethering (Fig.s 4C–D). Accordingly, at the low flow values studied here, both the chain cohesiveness and the interaction with the surface promoted the aggregation of tethered chains, of relevance for the VWF function (Fig.s 5B–C). Full phase diagrams had been previously determined for homo-polymers that adsorbed on homogeneous and non-homogeneous surfaces [33, 34]. Accordingly, our findings expand these studies by describing the complex interplay between chain-chain cohesion and chain-surface adhesion that influence the response of VWF to shear.

Self-association is a key feature for VWF function [20, 15, 16, 17, 23]. Self-association has the advantage of providing more adhesion sites to platelets, through indirectly immobilizing VWF chains [18]. Our results provide evidence of the spontaneous immobilization of self-associated VWF-like biopolymers (Fig.s 4–5). Our work quantifies the build-up of functionally absorbed aggregates in response flows at different cohesion/adsorption regimes and it thereby supports the notion of cooperative adsorption previously described for VWF [39].

Our work also compared the collective behavior of the polymer chains. Interestingly, for tethered chains at moderately-low flows, inter-chain interactions out-competed intra-chain ones resulting in a higher elongation of chains within an aggregate than in isolation (Fig.s. 3A–C). Furthermore, the elongation of tethered chains was similar to that of freely flowing bulk ones, except for chains that strongly interacted with the surface and thereby were less extended (Fig. 3D). Note that the term “tethered” conventionally refers to an end-attached polymer [20, 48]. In that case, tethered chains elongate more than bulk chains [49]. Here, we did not covalently immobilize the chains (as it has been done experimentally [22, 20]) but rather let them spontaneously bind to the surface, as it may be more likely to occur physiologically. A consequence of the multiple interaction points each VWF chain contains is that the strength of the interaction impacts the elongation of the chains deposited on the surface [41]. Hence, here we stress additional elongation properties that emerge when considering collectively multivalent adherent chains.

In our model, we adopted a CG resolution of one protein domain represented by one bead, as previously done to study D4-D4 interactions [40]. This level of resolution allowed us to take into account a more realistic distribution of adhesion points (corresponding to the VWF A3 domain). Surprisingly, the exposure did not significantly reduce when the interaction with the surface increased as it was observed for the chain elongation (Fig.s 6A,C). VWF has been commonly simulated at a lower level of resolution, typically one VWF monomer or dimer being represented by one or two beads [4, 33, 34, 40, 39, 35, 48, 38, 42]. When mapping our results to that low level of resolution, the surface area exposed to the solvent is much more sensitive to the interaction with the surface and correlates better with the trend exhibited by the chain elongation (Figure 6B,D). The exposed surface area is an essential functional quantity as it relates to the binding sites which are effectively available for the interaction of VWF with its partners, which are otherwise cryptic in the absence of shear. Conventionally, elongation is taken as a proxy for this observable. Our results confirm that qualitatively the correlation between elongation and surface exposure holds well, i.e. the more elongated the chains the more of the exposed area is available, but they more quantitatively coincide at a VWF-monomer size level of resolution. Ours constitutes an approach to connect fine-grain protein-domain details into coarser descriptions of VWF.

## Conclusion

Here, we studied the dynamics of polymers that resembled the blood protein VWF under shear flows by using Brownian dynamics simulations. Our simulations quantify the effect polymer-polymer cohesive and polymer-surface adhesion forces have on the flow-induced shear response of VWF. These forces coexist non-trivially to tune the elongation propensity of VWF and the formation of functional adsorbed VWF aggregates. Our data emphasizes the collective behavior of such aggregates featuring additional properties that VWF chains do not exhibit in isolation or freely flowing. Finally, we show that the resolution of the coarse-grain model impacts the exposed area of VWF aggregates. This systematic study is expected to contribute to our understanding of VWF and its ability to self-associate to accomplish its key hemostatic function.

## Author Contributions

H.A.-E. carried out the simulations. A.A.-K. and C.A.-S. designed the research. All authors analyzed the data and wrote the article.

## Acknowledgements

We thank Juan Carlos Briceño for the helpful discussions. For their funding, we are also grateful to the Max Planck Tandem initiative at the Universidad de los Andes (to H.A.-E and C.A.-S), the MIT International Science and Technology Initiatives (MISTI) Global Seed Funds (all authors), and to the Klaus Tschira Foundation (to C.A.-S.). We thank Frauke Gräter for the Ph.D. student internship (to H.A.-E.). We thank Guy Henderson for carefully reading the manuscript. We acknowledge the Max Planck Computing & Data Facility in Garching, Germany, and the HPC data center of the Universidad de los Andes in Bogota, Colombia, for the computing time and computational resources.

