## Supplementary information for "The interplay between adsorption and aggregation of von Willebrand factor chains in shear flows"

Supplementary Figures S1–S12.

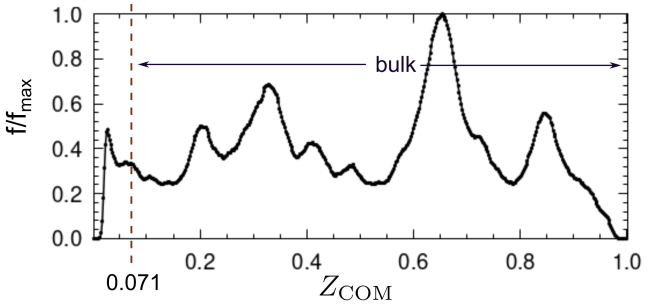


Figure S1: **Distribution f of the chain center-of-mass position along the coordinate orthogonal to the surface $Z_{\mathbf{COM}}$**. Distributions are normalized by the maximum observed value ($f_{max}$). To obtain this distribution all chain-chain cohesion energies ($\tilde{\epsilon}$=0.2, 0.4, 0.6, 0.8, and 1.0), chain-surface adhesion energies ($\tilde{\epsilon}_w = 2, 4$,6), and shear rates ($\tau\dot{\gamma}$ =0, 0.1, 0.25, 0.67, 1.55) were considered. The $Z_{\mathrm{cutoff}} = 7.12 \cdot 10^{-2}$, separating the bulk from the tethered region, is depicted with a dashed line.

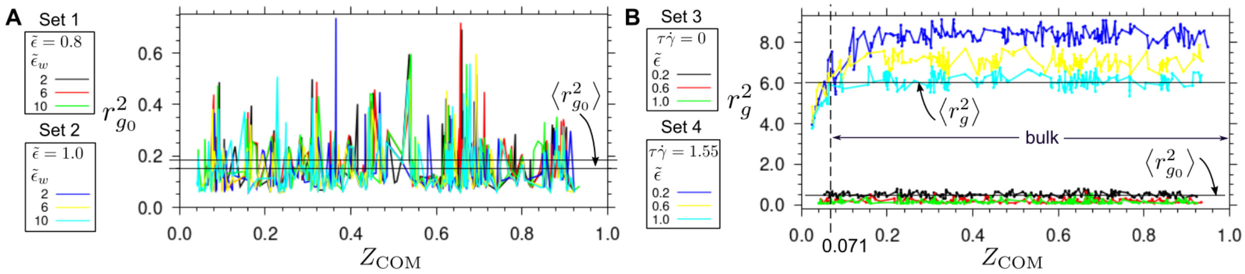


Figure S2: **Extension along $Z$ coordinate of VWF-like biopolymers**. (A) Normalized chain mean square radius of gyration ($r_g^2$) as a function of the chain average center of mass position $Z_{\mathrm{COM}}$. Every plot was obtained with the same chain-surface energy constant $\tilde{\epsilon}_w$ =10. Sets 1 and 2 represent plots with the same chain-chain interaction constant ($\tilde{\epsilon}$=0.8 and $\tilde{\epsilon}$=1.0, respectively) but different chain-surface one. Space between horizontal lines represents the range of average extension values $\langle r^2_{g_0} \rangle \sim 0.152 - 0.182$ for both sets. We obtained $\langle r^2_{g_0} \rangle$ ~0.169, 0.171, 0.182 and $\langle r^2_{g_0} \rangle$ ~0.152, 0.154, 0.165 for $\tilde{\epsilon} = 0.8$ and 1.0, respectively, corresponding to the adhesion constants $\tilde{\epsilon}_w$ in the same incremental order. (B) Normalized chain extension $r^2_{g_0}$ in the absence of flow, as a function of the chain $Z_{\mathrm{COM}}$. Sets 3 and 4 represent plots with the same flux ($\tau\dot{\gamma}$=0 and $\tau\dot{\gamma}$=1.55, respectively) but different cohesion energies. $r_g^2$ presented stable values along $Z$, with fluctuations about an average value $\langle r_g^2 \rangle$, when the chains were located in the bulk region ($Z > 7.12 \cdot 10^{-2}$). Horizontal black lines represent examples of average values per chain in the presence of flow $\langle r_g^2 \rangle$ or equilibrium $\langle r^2_{g_0} \rangle$.

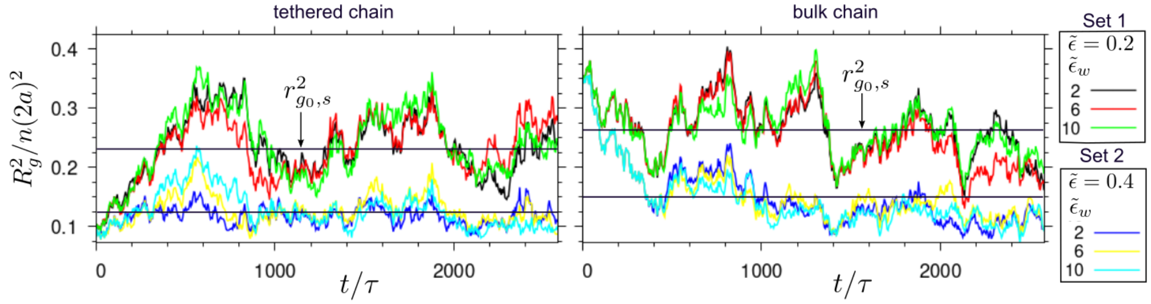


Figure S3: **Extension of a single VWF-like chain in the absence of flow as a function of time.** Extension is studied separately in the tethered and bulk regions. Sets 1 and 2 correspond to a chain-chain cohesion parameter of $\tilde{\epsilon}$=0.2 and $\tilde{\epsilon}$=0.4, respectively, varying the surface-chain adhesion energy (color lines). As a measure of the extension, the squared radius of gyration ($R_g^2$) normalized by the mean square extension of an ideal chain ($n(2a)^2$) is presented. Time is presented as a function of the characteristic time $\tau$. Note that the extension in each set yielded similar results independent of $\tilde{\epsilon}_w$. When time-averaging them, we obtained $r^2_{g_0,s}$ ~0.124 and 0.234 for the tethered chain, in sets 1 and 2, respectively, and $r^2_{g_0,s}$ ~0.150 and 0.264 for the bulk chain, in sets 1 and 2, respectively.

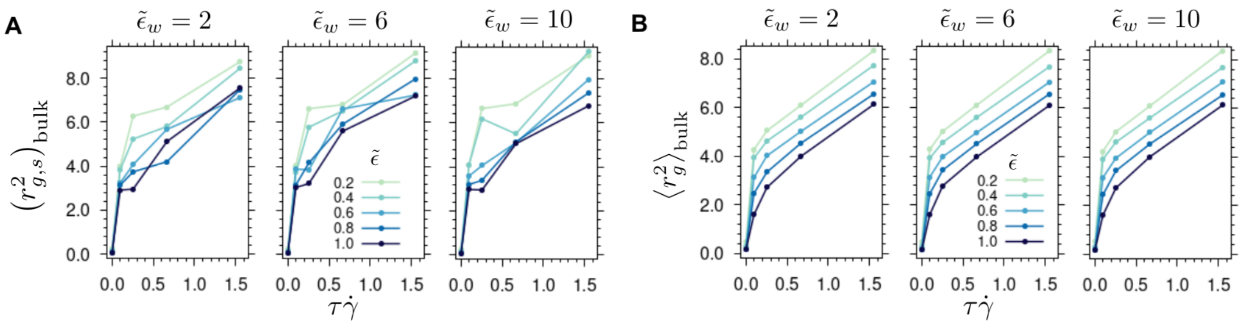


Figure S4: **Extension of bulk VWF-like biopolymers.** Bulk chain average extension as function of $\tau\dot{\gamma}$, per different $\tilde{\epsilon}$ (color) and $\tilde{\epsilon}_w$. (A) Normalized mean squared extension ($r^2_{g,s}$) for a single chain, flowing at the center of the box (bulk). (B) global mean square extensions $\langle r_g^2 \rangle$ for grouped bulk flowing chains.

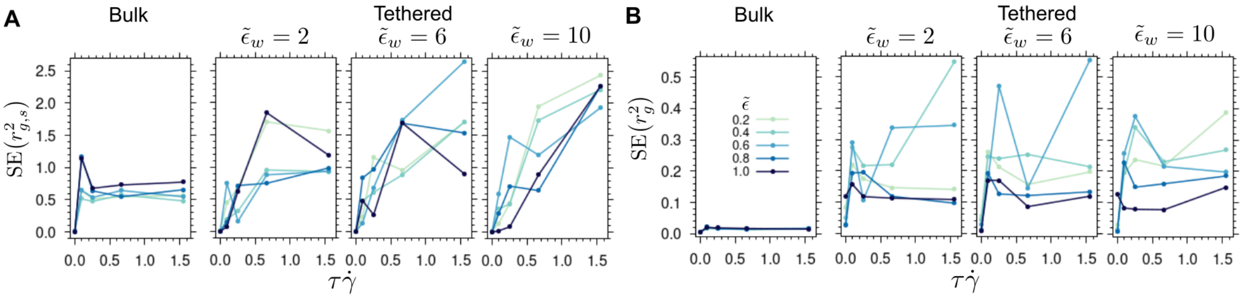


Figure S5: **Uncertainty in extension of VWF-like biopolymers**. Standard error (SE) in the extension of bulk and tethered chains, as a function of $\tau\dot{\gamma}$, for different $\tilde{\epsilon}$ (color) and $\tilde{\epsilon}_w$ values. (A) SE with respect to the mean $r^2_{g,s}$ for a single chain ($n_t \leq 158$ uncorrelated data samples were derived from the extension time series). (B) SE with respect to mean $\langle r_g^2 \rangle$ for the system with multiple chains ($n_s$ of independent samples here is the number of chains in either the bulk or the tethered region).

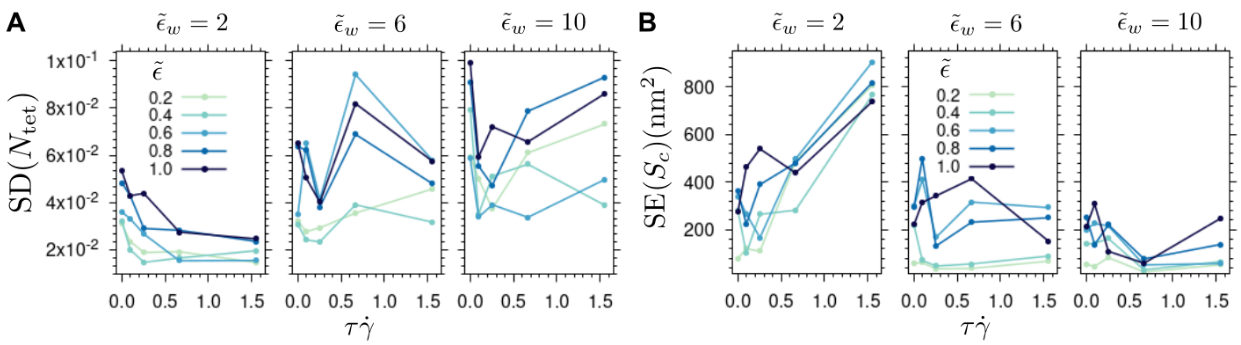

Figure S6: **Uncertainty in the number and exposure of tethered VWF-like biopolymers.** Standard deviation (SD) for the number $N_{\text{tet}}$ of tethered chains (A) and their normalized surface exposure (B) as a function of $\tau\dot{\gamma}$, $\tilde{\epsilon}$ and $\tilde{\epsilon}_w$. Uncertainty was estimated by bootstrapping in A (see methods). Up to $n_t = 196$ independent samples from the time series were obtained in B.

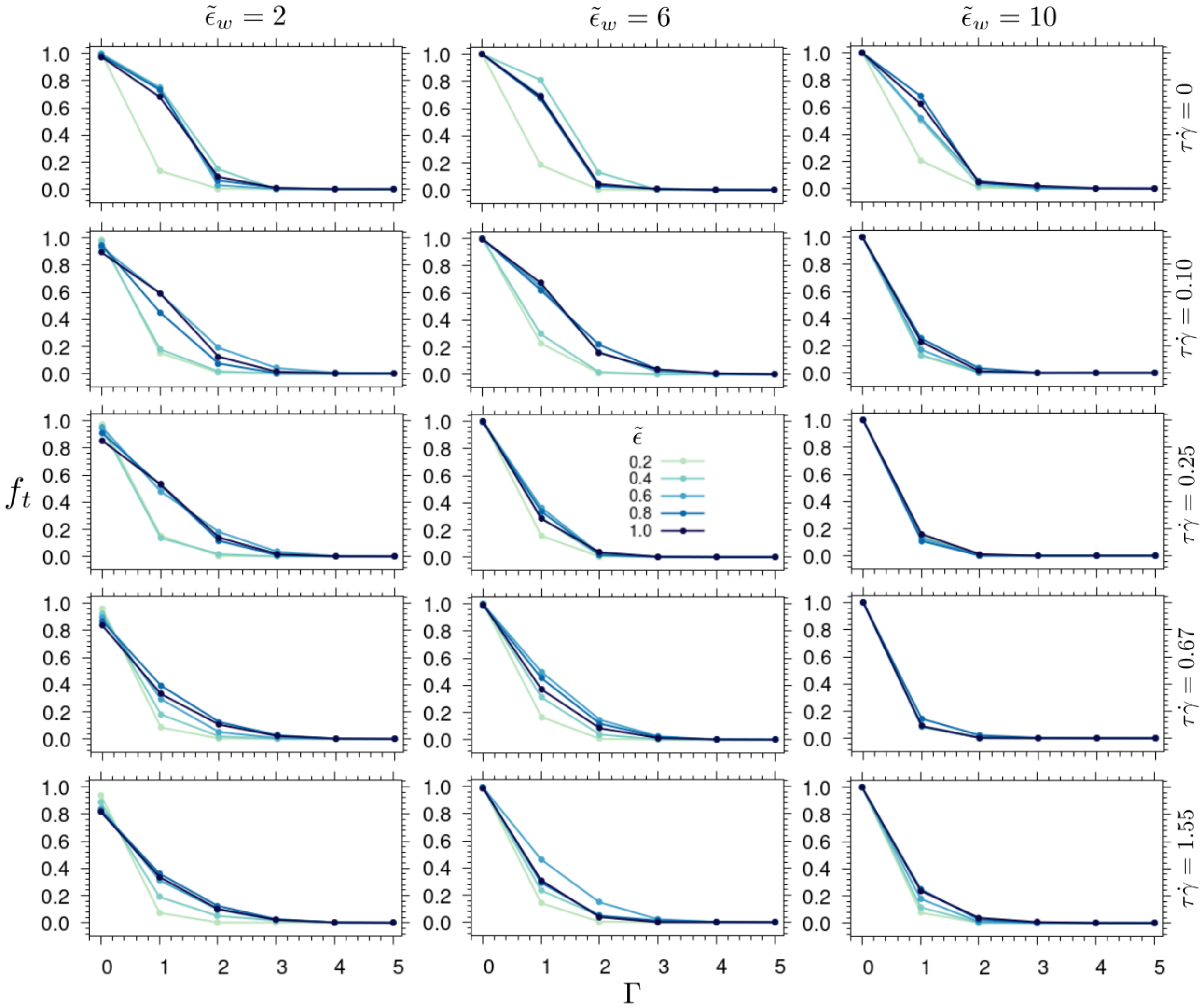

Figure S7: **Probability of having adjacent chains.** Fraction of time $f_t$ for which, one or more adjacent chains of type $\Gamma$ (see Fig. 4C) were observed, as a function of the flow (rows), adhesion (columns), and cohesion (color) energy. Mainly chains directly adsorbed ($\Gamma = 0$) were observed, then a lower probability of chains tethered to an adsorbed chain ($\Gamma = 1$). The probability for second neighbors or subsequent types ($\Gamma > 1$) increased with the augment of $\tilde{\epsilon}$, and decreased with the augment of $\tilde{\epsilon}_w$ and $\tau\dot{\gamma}$.

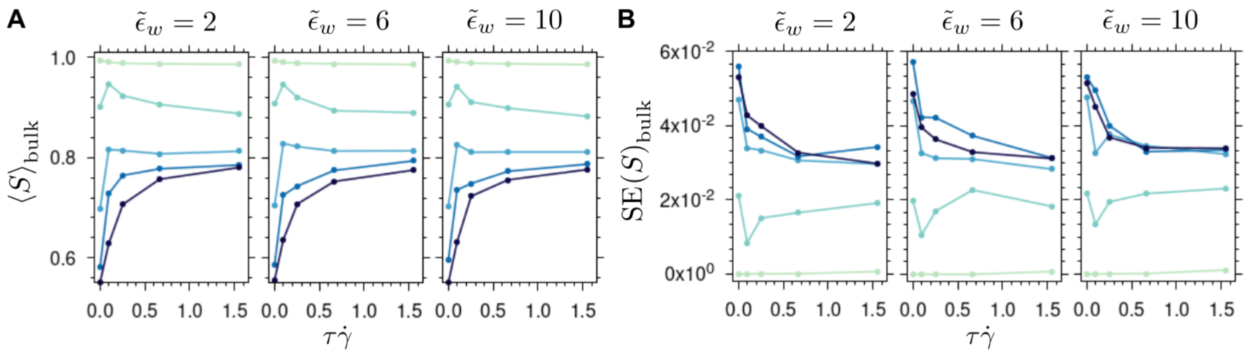


Figure S8: **Aggregation of bulk biopolymer VWF-like chains.** Time-averaged surface exposure ratio ($\langle S \rangle$) for bulk chains (A), and its error (B) are presented as a function of the shear rate ($\tau\dot{\gamma}$), and the cohesion $\tilde{\epsilon}$ (color) and surface $\tilde{\epsilon}_w$ (panels) energies.

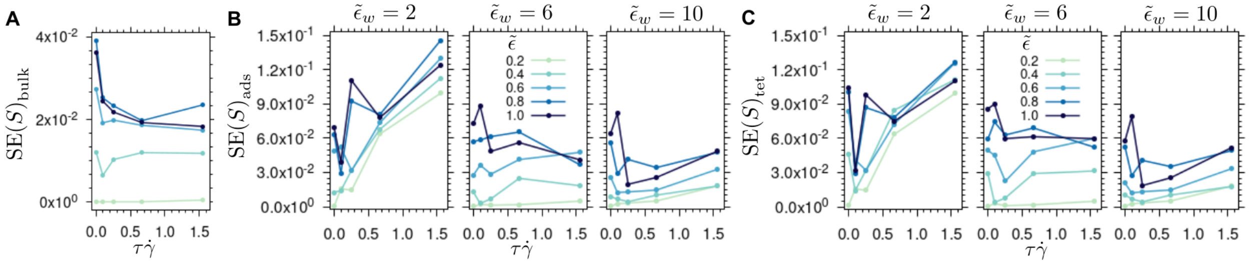


Figure S9: **Uncertainty in the aggregation of tethered VWF-like biopolymers.** Standard error (SE) in the exposure ratio ($\langle S \rangle$) is presented as function of the shear rate ($\tau\dot{\gamma}$), and the cohesion $\tilde{\epsilon}$ (color) and surface $\tilde{\epsilon}_w$ (panels) energies. SE was computed for bulk (A), adsorbed (B), and tethered chains (C). SEs computed in (A) take into account all the energies considered in Figure S8B. Up to $n_t$ =3333, 5000, and 769 uncorrelated data samples were obtained from the time series in A, B, and C, respectively.

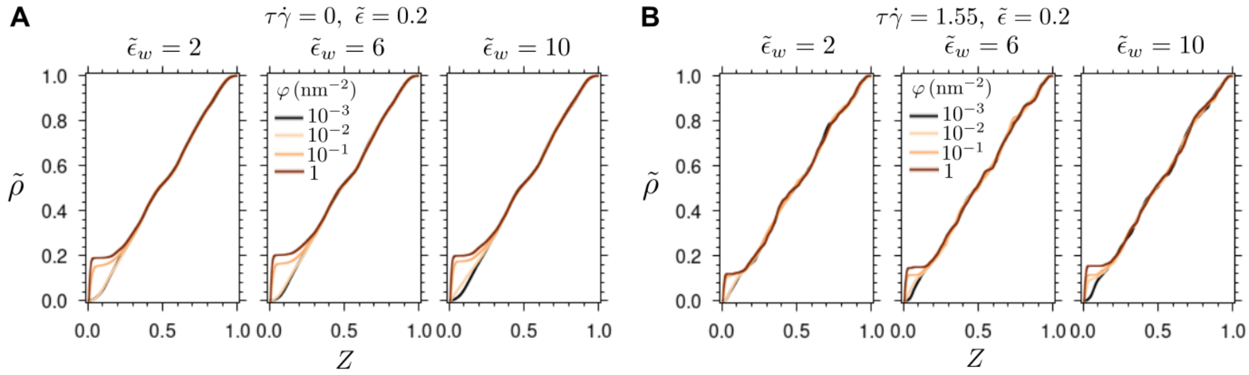


Figure S10: **Effect of density of receptors on the surface on the distance of the chains to the surface**. Normalized cumulative mass density $\tilde{\rho}$ along $Z$ axis (orthogonal to the surface) as a function of the density of receptors $\varphi$ and the chain–surface interaction strength $\tilde{\epsilon}_w$. Chain distribution was computed in the absence (A) or in the presence (B) of flow. A constant cohesion energy of $\tilde{\epsilon} = 0.2$ was considered here. For a high density of receptors ($\varphi = 10^{-1}$ and 1 receptors/nm$^2$) a first abrupt increment in $\tilde{\rho}$ near the adhesive surface ($Z = 0$), followed by constant values of $\tilde{\rho}$, indicates an accumulation of chains on the surface, but also an adjacent exclusion region with the absence of chains. For lower receptor densities ($\varphi = 10^{-3}$ and $10^{-2}$ receptors/nm$^2$) a constant augment of $\tilde{\rho}$ along $Z$ describes a homogeneous distribution of chains, in spite of the adhesion of the chain to the surface.

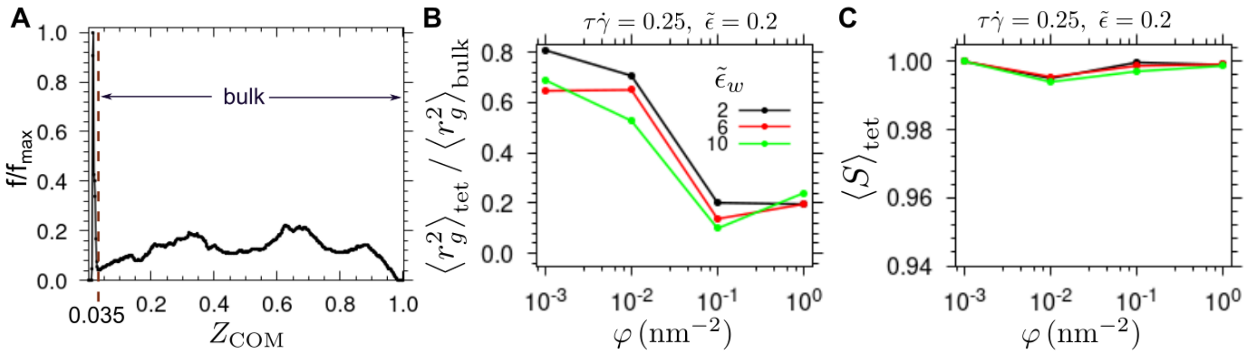


Figure S11: **Effect of surface density $\varphi$ of active sites on chain dynamics**. (A) Distribution f of the chain center-of-mass position along the coordinate orthogonal to the surface $Z_{\mathrm{COM}}$. Distributions are normalized by the maximum observed value ($f_{max}$). Here the distribution was obtained considering systems with different surface densities ($\varphi = 10^{-3}, 10^{-2}, 10^{-1}, 1$ nm$^{-2}$), adhesion constants ($\tilde{\epsilon}_w = 2, 4$,6), and shear rates ($\tau\dot{\gamma}$ =0, 0.25, and 1.55). (B–C) Extension ratio of tethered to bulk flowing chains (B) and time-averaged surface exposure ratio of tethered chains (C) as a function of the receptor density $\varphi$, for the indicated cohesion energy and shear rate, but variable adhesion constant (color). The cutoff separating both regions was $Z_{\mathrm{cutoff}} = 0.035$, according to the end of the peak near the surface (see A).

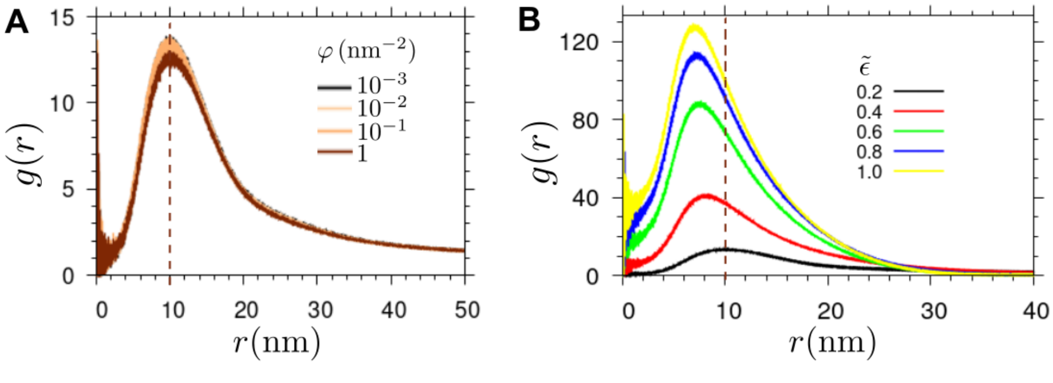


Figure S12: **Radial distribution function** ($g(r)$) of the center of mass of the VWF monomers recovered from the protein-domain BD trajectories. $g(r)$ is shown as a function of the surface density $\varphi$ averaging over multiple cohesion energies (A) and the cohesion energy $\tilde{\epsilon}$ for a constant $\varphi = 10^{-2}$ (B). A VWF monomer was assumed to have a size of $r = 10$ nm (dashed line).
